# Ketosynthase engineering enhances activity and shifts specificity towards non-native extender units in type I linear polyketide synthase

**DOI:** 10.1101/2023.02.16.528826

**Authors:** Patrick D. Gerlinger, Georgia Angelidou, Nicole Paczia, Tobias J. Erb

## Abstract

Engineering modular type I polyketide synthases (PKS) for the targeted incorporation of non-natural substrates to create variations in the polyketide backbone is a long-standing goal of PKS research. Thus far, most approaches focused on engineering the acyltransferase domain (AT) of PKS, whereas the effects of other ubiquitous domains such as the ketosynthase domain (KS) have received much less attention. In this work, we investigated the effects of thirteen active site substitutions in the module 3 KS (KS3) of the 6-deoxyerythronolide B synthase (DEBS) on incorporation of non-natural extender units *in vitro*. Using a truncated and a complete DEBS assembly line, we show that substitutions of F263 in KS3 invert specificity up to 1,250-fold towards incorporation of non-natural extender units in the terminal position. In contrast, substitutions of I444 in KS3 show up to 8-fold increased production of 6-deoxyerythonolide B (6-dEB) analogues with non-natural extender units at internal positions. The latter notably without compromising overall productivity of the assembly line. Our study further elucidates the underlying mechanisms for these different behaviors, highlighting the potential of KS engineering for the production of designer polyketides in the future.

## Introduction

Type I Polyketide Synthases (PKSs) are multi-domain enzyme complexes that produce bioactive secondary metabolites with remarkable structural and functional diversity, including antibiotics, antifungals and anticancer compounds^1-3^. The chemical backbone of all these compounds is usually derived from a priming acyl-CoA starter unit and a simple set of (alkyl)malonyl-CoA extender units, such as malonyl-, methylmalonyl-, or ethylmalonyl-CoA, which are condensed and further modified through the action of different domains on the growing polyketide chain^4,5^.

Type I PKSs are organized within modules that each consist of at least three core domains – a ketosynthase (KS), acyltransferase (AT) and an acyl-carrier protein (ACP) that together select the extender unit substrate and catalyze a decarboxylative Claisen condensation reaction for chain elongation. Additional domains, including ketoreductases (KR), dehydratases (DH) and enoylreductases (ER) can create further chemical diversity through epimerizations of the α-alkyl-residue, and/or the introduction of β-hydroxyl groups, α,β-desaturated or fully saturated α,β-bonds in the growing polyketide chain.

The arguably most well studied model system for type I PKS is DEBS, the 6-deoxyerythronolide B synthase. DEBS consists of six modules, each of which catalyze one round of chain elongation through the incorporation of methylmalonyl-CoA. The assembly line is primed with propionyl-CoA through an N-terminal loading domain, and terminated via cyclization through a thioesterase, producing 6-deoxyerythronolide B (6-dEB), the macrolide backbone of the antibiotic erythromycin.

In linear type I PKS, such as DEBS, the sequence of reactions, as well as substrate- and intermediate-specificity are encoded in conserved fingerprint motifs within the different domains^6,7^, a concept commonly referred to as co-linearity^8,9^. This design principle of type I PKS assembly lines has raised the vision of generating designer polyketides through domain re-programming, either via mutagenesis approaches or replacements of individual domains and/or modules. These strategies aimed at changing the redox state of the polyketide (e.g., by engineering or transplanting KR and ER domains^10,11^), or enabling the incorporation of non-native and non-natural malonyl-CoA analogues to modify the structural backbone of the polyketide. The latter has been mainly attempted by engineering of the substrate specificity of AT domains. While AT domain engineering was met with some success in the past ^12-18^, these efforts were mainly successful when focused on engineering a terminal or standalone PKS module (see below), which has led to the realization that AT engineering (still) has its limitations.

Contrary to the AT, KS domains have received little attention in respect to specificity-engineering approaches, although several studies hint towards KS domains as limiting factor for the incorporation and processing of non-native and non-natural substrates in the growing polyketide chain. First, KS domains were shown to be the rate-limiting step in polyketide chain elongation^19-21^, which could be overcome by single amino acid substitution^22^. Second, it was shown that KS domains exert a gatekeeping function, which allowed the identification of different sequence motifs corresponding to substrate specificities^23,24^. Finally, it has been realized lately that the downstream KS domains co-migrate with their cognate upstream module, ^25,26^, which has led to a redefinition of the boundaries of PKS modules and identified KS domains as interesting target for specificity engineering efforts.

Here, we explored the effect of single active-site residue substitutions in the KS of DEBS module 3 in respect to the incorporation of non-native alkylmalonyl-CoAs *in vitro*. We identified seven potential target sites through a sequence-structural modelling approach and evaluated a total of 13 different mutations. We show that one set of mutations increases incorporation of non-native extender units into the terminal position of a DEBS tetraketide system with specificity inversion of up to 1,250-fold. We further show that another set of mutations lead to up to 8-fold increased incorporation of non-native extender units in the internal position of a full DEBS assembly line. Notably, and in contrast to many other efforts, the latter does not come at the cost of decreased productivity of the DEBS PKS, truly improving modified polyketide yield. Overall, our study demonstrates that KS engineering is a so far underexplored, but promising strategy towards the production of modified polyketides in the future.

## Main

To study the effect of KS engineering onto incorporation of non-natural extender units, we focused our attention on DEBS and in particular the AT domain of module 2 (AT2). AT2 of DEBS is naturally promiscuous and has been reported to incorporate longer chain alkylmalonyl-CoAs beyond its natural substrate methylmalonyl-CoA (Figure 1, compound **a**)^27,28^. Thus, we decided to test whether active site mutations of KS of module 3 (KS3) would increase the incorporation of longer alkyl-chains by DEBS module 2.

**Figure 1.**
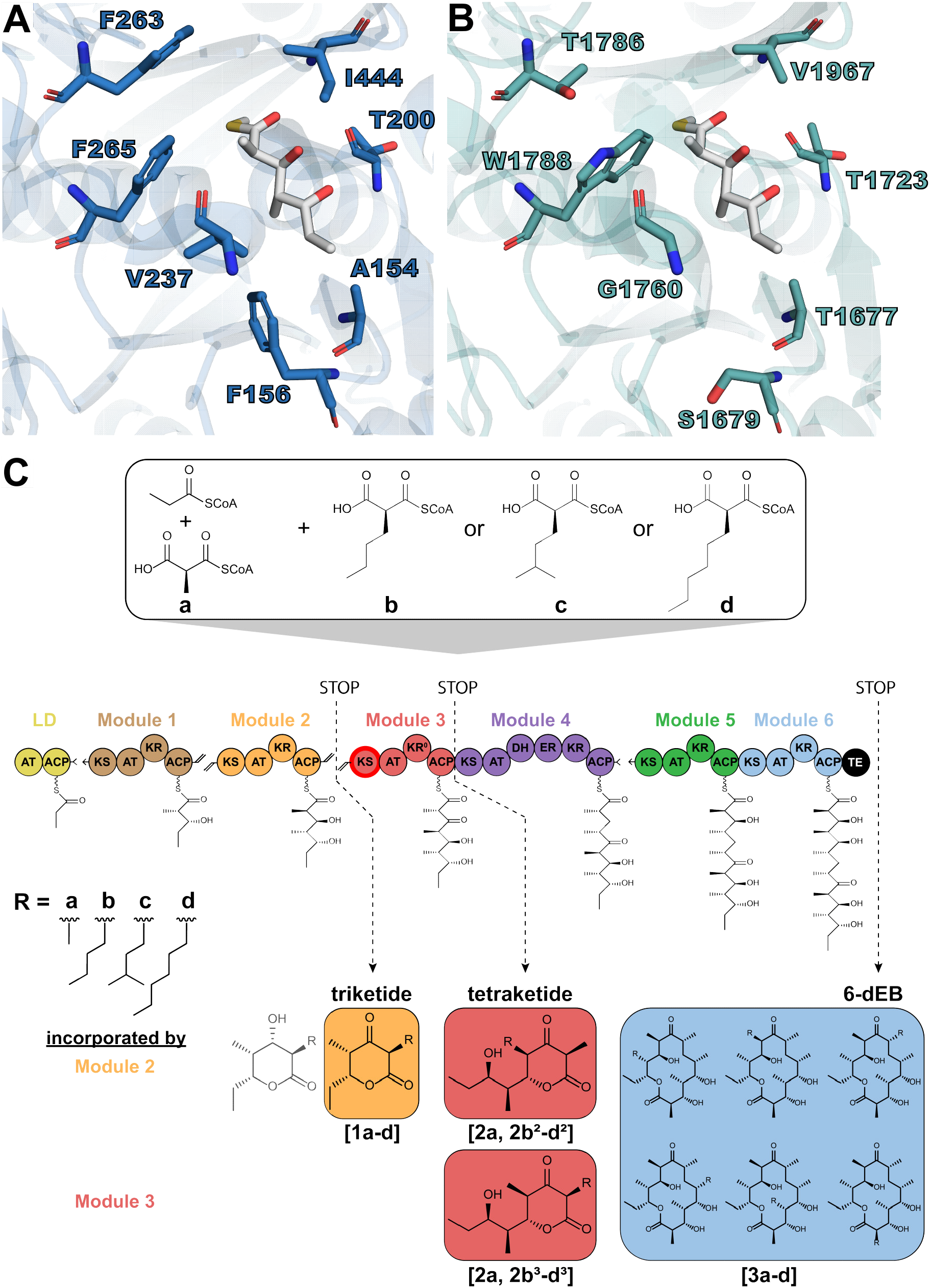
Assay setup, detected polyketides and structural comparison for engineering strategy. **A** Residues of the KS3 active site (PDB: 2QO3)^29^ chosen for mutagenesis, as well as the corresponding residues in a homology model of the reveromycin KS5 active site (**B**). Additionally displayed is a triketide intermediate, the α-methyl-moiety pointing upwards towards F265 and I444 in **A** (adapted from Hirsch et al.)^24^. Residue numbering is according to the crystal structure for KS3, and adapted for Rev KS5 using the protein residue numbering of revB (as given by the ClusterCAD database)^32^. **C** Each assay contains propionyl-CoA, methylmalonyl-CoA (a) and either non-natural extender unit butylmalonyl-CoA (b), 3-methylbutylmalonyl-CoA (c) or hexylmalonyl-CoA (d). For the exact assay setup, see Methods. DEBS had been reconstituted according to literature^28^, and would produce each shown product (triketide **1**, tetraketide **2**, and 6-dEB heptaketide **3**). The truncated system lacked downstream modules after module 3 (tetraketide system), and would produce all products up until the termination point. Products are labeled according to the termination points and the tetraketide additionally distinguished regarding side-chains from extender unit incorporation by modules 2 or 3 (denoted as superscript). Note that an incorporation of **b-d** by module 1 was observed in the triketide, but not in the tetraketide, and is therefore not displayed for simplicity. Potential 6-dEB isomers could not be separated and are therefore summarized without superscript labeling.

To identify potential sites for active site engineering, we performed a sequence-function based alignment of 600 reviewed KS sequences to assess the conservation of active site residues in proximity of the conserved catalytic triad (Figure S1)^29^. We also used information from a recent study that was able to correlate certain KS active site residues to size and configuration of nascent the polyketide chain^24^.

To further validate these target sites, we compared the crystal structure of the DEBS KS3-AT3 didomain^29^ and a homology model of the reveromycin KS5 domain^30^. The reveromycin KS5 domain processes larger α-alkyl substituents (C4-C6) but otherwise shares an identical configuration of the α- and β-substituents with DEBS KS3. Our analysis revealed a hydrophobic area in DEBS KS3 comprised of F263, F265 plus V237, and I444, which together restrict the active site area and likely exclude larger alkyl-substituents (Figure 1A). In reveromycin KS5, on the other hand, residues T1786 (corresponding to F263), W1788 plus G1760 (corresponding to F265 plus V237) and V1967 (corresponding to I444), respectively form a larger cavity that is upwards oriented and able to accommodate the natural C4-6 alkyl chain during biosynthesis^30^ (Figure 1B).

Notably, some of the KS3 residues of DEBS that we identified (V237, F263 and F265) plus others that are in close proximity to the polyketide chain (F156, T200) or the KS-KS dimerization surface (A154) had previously been targeted in different engineering efforts^19,22,31^, some of which had resulted in increased turnover rates. However, the effects on promiscuity towards non-native substrates in these mutants had not been systematically tested, with exception of A154W and F263T, which showed broadened substrate specificity towards a variety of *N*-acetylcysteamine thioester substrates^22^. Thus, we decided to target those seven active site residues and test their effects on the incorporation of four different non-natural extender units (Figure 1 compound **b**, butylmalonyl-CoA; **c**, 3-methylbutylmalonyl-CoA; **d** hexylmalonyl-CoA) in a tetraketide DEBS (“truncated assembly line”) and a hepatketide DEBS (“full assembly line”) system.

To assess the effects on polyketide product formation, we used a qualitative HPLC-MS based method. This enabled us to compare product formation (tetra-or heptaketide) of the respective wildtype (WT) DEBS system with the different DEBS variants. Besides monitoring tetra-or heptaketide product formation, our HPLC-MS method also allowed us to observe (shunt) products of the respective assembly lines (Figure 1C) to monitor premature release of the growing polyketide chain.

We introduced a total of 13 different substitutions at the seven target sites in the DEBS tetraketide system and probed for incorporation of extender units **b-d** into the expected tetraketide product. LC-MS analysis showed two baseline-separated product peaks with identical masses (Figure S2). Tandem MS analysis revealed those to be tetraketide isomers, carrying alkyl-substitutions at either the C4 or C2 position (see Methods); the former (C4 position) corresponding to internal incorporation of **b-d** by module 2 (**2b**^**2**^**-d**^**2**^), the latter (C2 position) corresponding to terminal incorporation of **b-d** by module 3 (**2b**^**3**^**-d**^**3**^). Note that we did not detect products corresponding to multiple incorporations of **b-d**, or a significant increase of triketides **1b-d**, indicating premature termination product due to decreased processing by module 3 (data not shown).

While we observed no significant changes for variants A154W, T200A or I444V and I444A, both V237G and V237A variants showed a 4-to 5-fold increase of C4-substituted tetraketide **2d**^**2**^, indicating an enhanced tolerance of KS3 towards a hexyl-substituted intermediate from incorporation of **d** by module 2. These results were in line with the hypothesis that KS domain engineering increases processing of non-natural extender units that are incorporated in an internal position through the AT of the upstream module.

Intriguingly, all variants of F263 (F263T and F263A), as well as all variants of F265 (F265A, F265M and F265V) displayed an increase in the C2-substituted tetraketide, corresponding to an increased incorporation of non-natural extender units in the terminal position (**2b**^**3**^**-d**^**3**^, Figure 2A). The effect was more pronounced for the F263 variants, with F263T producing 24-, 55- and 25-fold more **2b**^**3**^, **2c**^**3**^ and **2d**^**3**^, respectively. The F263A variant even exceeded those yields with a 26-, 75-, and 48-fold increase in formation of products **2b**^**3**^, **2c**^**3**^ and **2d**^**3**^. Notably, this increase in C2-substituted products came at the expense of natural tetraketide **2a** (Figure 2B). Product **2a** decreased around 10-to 20-fold compared to the WT in assays containing **a** and either **b, c** or **d**. In combination with the 20-70-fold increased incorporation of **b-d** at position C2, this implies a specificity change between more than 100-fold towards **d** (hexylmalonyl-CoA) by F263T and up to 1,250-fold for **c** (3-methylbutylmalonyl-CoA) by F263A (Table 1). This change was also reflected by an increase in the percentage yield of non-natural tetraketide product (Table 2). Compare to the WT that produced less than 5% of either **2b³**-**d³**, relative yields increased up to 90% for F263T and over 90% for **2b³-d³** for F263A.

**Figure 2.**
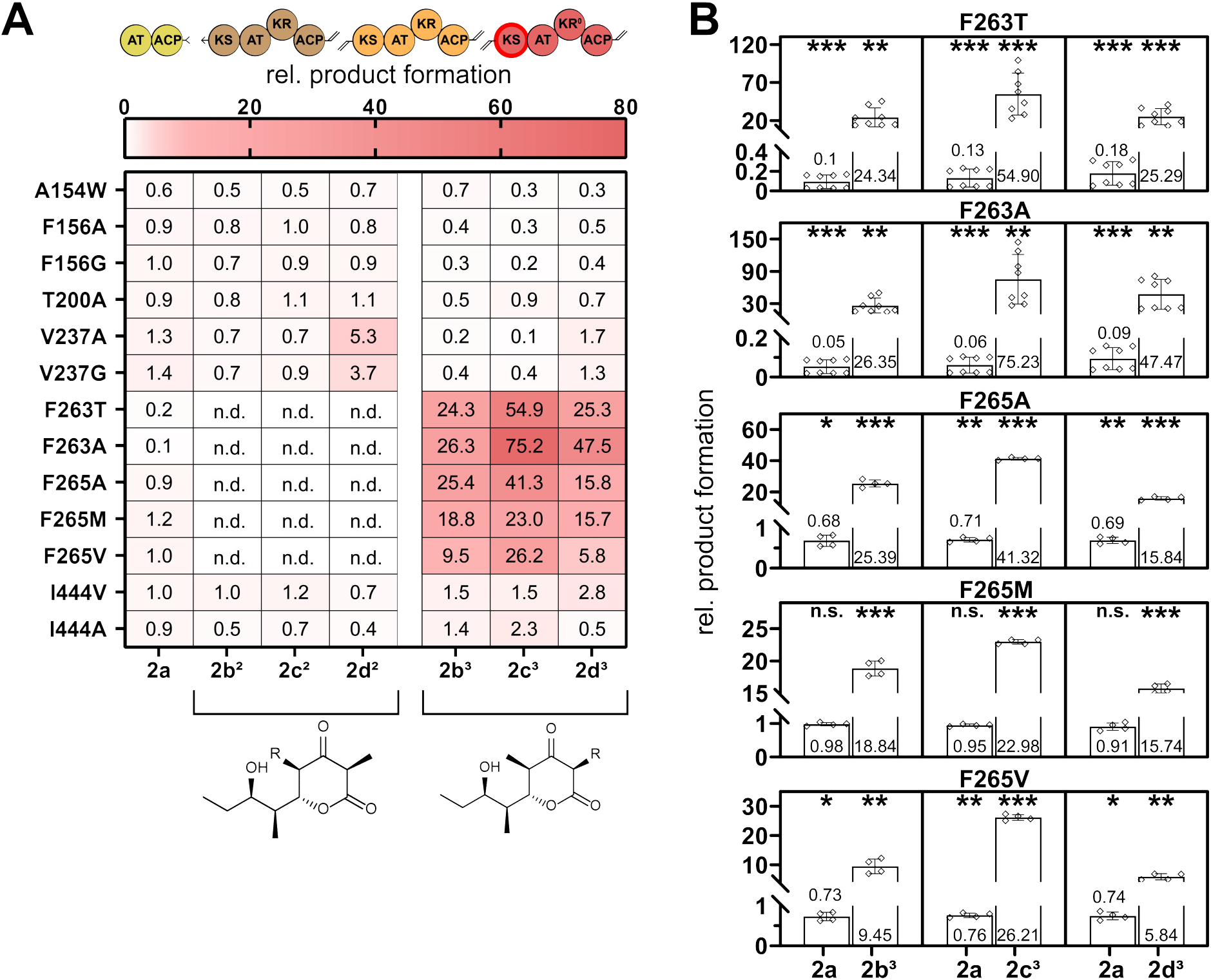
Relative product formation of the KS3 variants. **A** tetraketide analogues produced by KS variants in the tetraketide system. **B** production of 2a in tetraketide assays containing non-natural substrates, together with the product formation from module 3 incorporation as in A. Color usage is according to Figure 1. Values are derived from the detected EIC of each compound compared to that of the wild type after 17 h and display the mean of *n* = 2 biological replicates (each in duplicates; two technical replicates for F265 variants). Significance (H0: relative product formation = 1) was determined using a two-tailed one-sample t-test. *p ≤ 0.05, **p ≤ 0.005, ***p ≤ 0.0005. n.d. – not detected, n.s. – not significant.

**Table 1.** Apparent specificity changes. Values are given as product ratios of non-natural tetraketide (**2b³, 2c³** or **2d³**) over natural tetraketide (**2a**) from the data displayed in Figure 2B.

|  | x-fold shift towards non-natural tetraketide |  |  |
| --- | --- | --- | --- |
|  | **b** | **c** | **d** |
| **F263T** | 243 | 422 | 141 |
| **F263A** | 527 | 1254 | 527 |
| **F265A** | 37 | 58 | 23 |
| **F265M** | 19 | 24 | 17 |
| **F265V** | 13 | 34 | 8 |

**Table 2.** Percentage of b-d derived tetraketides. Values were taken from the respective EICs summarized for tetraketide produced by the F263 variants. Displayed is the mean ± standard error.

|  | % product |  |  |
| --- | --- | --- | --- |
|  | **2b<sup>3</sup>** | **2c<sup>3</sup>** | **2d<sup>3</sup>** |
| **WT** | 4 ± 1 | 2.8 ± 0.7 | 3.0 ± 0.7 |
| **F263T** | 89 ± 4 | 89 ± 3 | 76 ± 5 |
| **F263A** | 94 ± 2 | 95.9 ± 0.6 | 90.8 ± 0.6 |

**Table 3.**
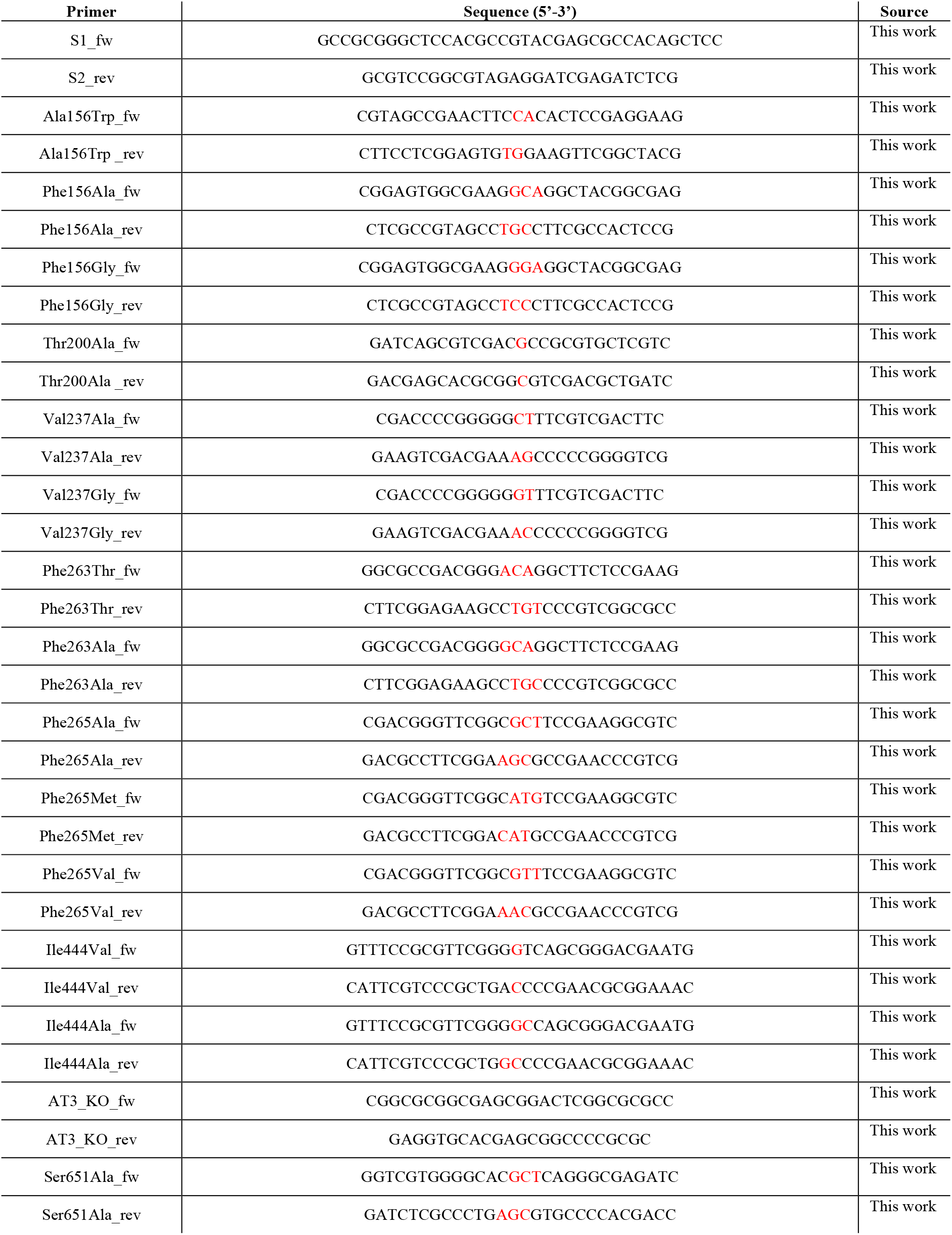
All primers used in this work. Mutations are depicted in red.

**Table 4.** Buffers used in this work.

|  |  |
| --- | --- |
| Buffer A | 450 mM NaCl<br>15 % glycerol<br>50 mM phosphate<br>pH 7.6 |
| Buffer B | 50 mM NaCl<br>10 % glycerol<br>50 mM phosphate<br>500 mM imidazole<br>pH 7.6 |
| Buffer C | 10 % glycerol<br>50 mM phosphate<br>pH 7.6 |
| Buffer D | 500 mM NaCl<br>20 % glycerol<br>50 mM phosphate<br>pH 7.6 |
| Buffer E | 350 mM NaCl<br>10 % glycerol<br>50 mM phosphate<br>pH 7.6 |
| Buffer F | 450 mM NaCl<br>15 % glycerol<br>50 mM HEPES<br>pH 7.6 |
| Buffer G | 50 mM NaCl<br>500 mM imidazole<br>10 % glycerol<br>50 mM HEPES<br>pH 7.6 |
| Buffer H | 350 mM NaCl<br>20 % glycerol<br>50 mM HEPES<br>pH 7.6 |

Similar to the F263 variants, all three F265 variants (F265A, F265M and F265V) also showed a strong increase in **b-d** derived tetraketide product (Figure 2B), albeit to a slightly lesser extent than the F263 variants (between 6-to 42-fold). Formation of **2a** was also less affected (approximately a 1.5-fold decrease for both F265A and F265V, no significant difference in case of F265M). Overall, the F265 variants displayed a change in specificity towards the respective non-native extender unit between 8- and 60-fold (with the largest shifts in specificity towards **c**, Table 1). Together, these results for the F263 and F265 variant indicated a specificity inversion in the terminal position of the tetraketide DEBS system, although they did not provide a mechanistic explanation, yet (see below).

To assess how above findings would translate in respect to the production of 6-dEB analogues, we tested the 13 different KS3 substitutions also in the context of the complete DEBS assembly line. Unfortunately, the F263 variants, which had shown a specificity switch at the terminal position in the tetraketide system (see above) did neither produce 6-dEB (**3a**) nor non-natural analogues (**3c-d**) in the presence of non-native extender units. Moreover, formation of 6-dEB was decreased between 5-to 10-fold in assays containing only methylmalonyl-CoA (Figure 3A). However, we observed a 3-fold increase of triketide product **1c** in the F263 variants (Figure 3A). Furthermore, the F263 variants showed increased production of tetraketides **2b**^**3**^**-d**^**3**^ with inverted specificities (between 10- and 60-fold, Figure 3A, 3B), indicating that the F263 mutations had created a path towards formation of dead-end shunt products at the level of tetraketides.

**Figure 3.**
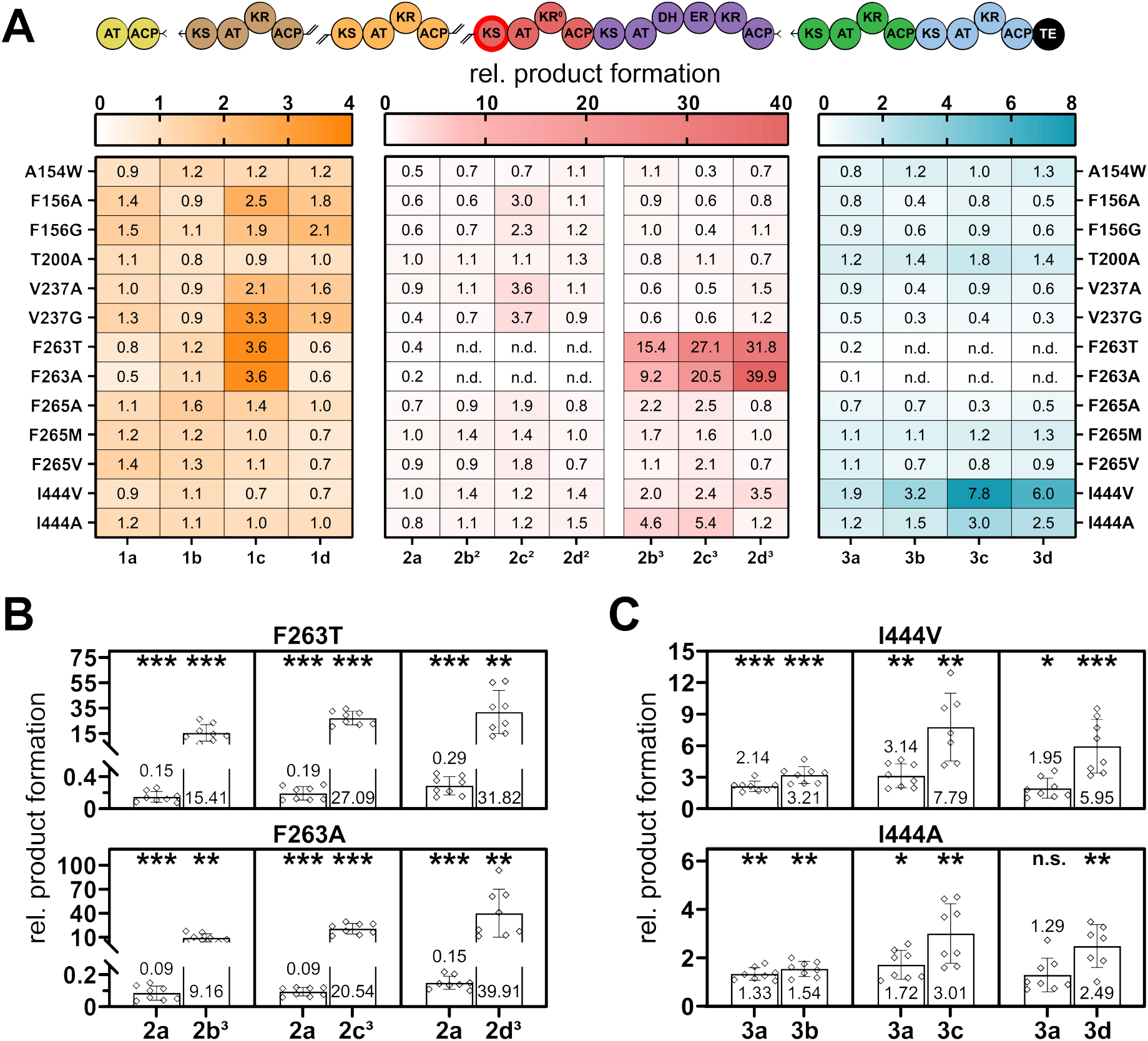
Relative product formation of the KS3 variants. **A** Product formation pattern produced by KS variants in the full DEBS system. **B** Production of 2a in DEBS assays containing non-natural substrates, together with the product formation from module 3 incorporation as in A. **C** Product formation of 3a in DEBS assays containing non-natural substrates, together with the product formation of **3b-d** as in A. Color usage is according to Figure 1. Values are derived from the detected EIC of each compound compared to that of the wild type after 17 h and display the mean of *n* = 2 biological replicates (each in duplicates; two technical replicates for A154W, F156A, T200A, V237A and V237G variants). Significance (H0: relative product formation = 1) was determined using a two-tailed one-sample t-test. *p ≤ 0.05, **p ≤ 0.005, ***p ≤ 0.0005. n.d. – not detected. n.s. – not significant.

We hypothesized that this behavior of the F263 variants was the result of an AT3-ACP3 bypassing mechanism as proposed by Ad et al. ^33^ (see Figure S3 and Discussion). To validate this hypothesis, we created a catalytic knockout of AT3 (S651A). When testing the S651A variant alone, relative product formation decreased to 10-40%, while in combination with the F263T variant, double variant S651A F263T, showed product yields similar to the F263T variant alone (Figure S4A, S4B). This demonstrated that incorporation of non-natural extender units in the F263 variants was independent of AT3 activity, supporting the hypothesis of an intramodular AT-ACP bypassing mechanism. To exclude that the effect might stem from substrate competition, we additionally tested the F263 variants with 10-fold lower concentration of **b-d** (100 µM instead of 1 mM), but did not observe any changes (Figure S4A, S4C), supporting the AT3-ACP3 bypassing hypothesis for these mutations.

In contrast to the F263 variants, all three F265 variants that also had shown a strong specificity switch in the terminal position of the tetraketide system, displayed product patterns similar to the WT in the complete assembly line, with no significant increase of shunt products or 6-dEB analogues. The only effect observed was a decrease of **3a-d** for the F265A variant (Figure 3A, Figure S5). In summary, this data indicated that the AT3-ACP3 bypass mechanism was not pronounced in the full assembly line for the F265 mutants.

Much to our surprise, I444 variants I444V and I444A, which had not shown any effects in the tetraketide system, resulted in a significant increase of **b-d** in the final product, when tested in the complete assembly line, and only formed some shunt products **2b**^**3**^-**2d**^**3**^ (Figure 3A). Formation of products **3b-d** was increased two-to threefold for I444A, and between two-to eightfold for I444V, while the production of **3a** remained constant or increased up to threefold in the presence of non-natural substrates (Figure 3C). In case of the I444V variant, the product profile was shifted towards non-natural products (Figure 3C). To the best of our knowledge, this is the first successful engineering attempt of a KS in a complete polyketide assembly line and even more notable, as it does not come at the expense of a reduced productivity compared to the wild-type.

Although we could not determine the exact position of incorporation in the 6-dEB the shift in the product spectrum of the I444V variant strongly suggest that incorporation is site specific, but might still be suffering from (downstream) kinetic bottlenecks. This was further supported by findings that efforts to increase incorporation of non-native extender units through additional AT2 engineering did not further increase productivity of the overall system (see Supplementary Information, Figure S6-S8). Nevertheless, these results demonstrated that KS3 engineering is a promising strategy to increase incorporation of non-native extender units at internal positions in a full type I PKS assembly line without compromising overall productivity.

## Discussion

In this work, we studied the impact of different active site mutations onto the incorporation of non-natural extender units in a truncated, as well as a complete DEBS PKS assembly line. We observed different outcomes depending on the mutation site, as well as the incorporation site (internal versus terminal). These effects can be explained by different underlying mechanisms that might be exploited for PKS engineering in the future.

Variants of F263 (and F265) showed an almost complete inversion in specificity from methylmalonyl-to longer-chained alkylmalonyl-CoA in the terminal position of the truncated DEBS system. However, this comes at the expense of a shunting mechanism which terminates the assembly line at module 3 (Figure 3A, Figure S3). F263 confines the active site entrance boundaries of KS3 for the (methyl)malonyl-ACP3 reaction partner. This constraint is relieved by introducing a threonine or alanine, likely allowing longer chain alkylmalonyl-CoAs to diffuse into the KS3 active site, where they can function as nucleophile instead of the ACP-substrate.

This reaction does not require a conformational change of the KS and is in line with the proposed “turnstile mechanism”^34,35^, ultimately resulting in a CoA-tethered product. This CoA-bound polyketide poses a true dead-end shunt product and can further undergo cyclization, releasing the corresponding tetraketide. At the same time opening the active site entrance probably also negatively affect interaction and/or positioning of the original substrate, thus lowering the rate with the original substrate methylmalonyl-ACP/CoA. While the shunting mechanism can currently be “only” exploited for the incorporation of non-native extender units at the terminal position, it might be leveraged in context of the complete assembly line by providing extender units bound to a *trans*-ACP, instead of CoA, to promote downstream processing. Such ACP-bound extender could be constantly recycled through an ACP-thioester regenerating system and eventually allow exploitation for site-specific incorporation at internal positions in the future ^36^.

In contrast, I444 variants, which did not show increased any effects in the tetraketide model system, displayed up to 8-fold increased incorporation of non-natural extender units for the production of 6-dEB analogs (**3b-d**), with some accumulation of (shunt) products (**2b**^**2**^**-d**^**2**^) but notably without compromising overall productivity of the wild-type system. This strongly suggested that the I444 mutation increased promiscuity of the KS3 towards processing of non-natural extender units incorporated by upstream modules. Thus, the I444 variants represent a step forward for efforts for the site-specific incorporation of non-natural extender units at internal positions.

Taken together, our findings highlight the potential of KS domains as engineering target for the incorporation of non-natural moieties at different positions into the polyketide product backbone. Notably, some of the amino acids investigated in this study (e.g., F265 and I444) show a high degree of conservation among type I KS domains (Figure S1). It will be interesting to assess the effect of mutagenesis of these residues in other PKS systems to explore and eventually exploit KS domain engineering as strategy for the production of modified polyketides in the future.

## Methods

Chemicals were obtained from Sigma-Aldrich and CARL ROTH, unless otherwise denoted. Coenzyme A was purchased from Roche Diagnostics. Primers were purchased from Eurofins MWG. Materials and equipment for protein purification were obtained from Cytiva.

Q5 HotStart Polymerase (New England Biolabs) was used according to protocol (including high GC buffer) for all PCR amplifications. Gibson assembly was performed using Gibson Assembly MasterMix (New England Biolabs), a backbone/insert ratio of 1:3 (20 pmol backbone DNA) and an incubation time of 40 min at 50°C. Ligation was carried out according to protocol for the T4 ligase (New England Biolabs) with a backbone/insert ratio of 1:3 (20 pmol backbone DNA) and an incubation time of 20 min at RT. For restriction enzyme digest of DNA, FastDigest Enzymes (Thermo Fisher Scientific) were utilized according to recommended incubation times and temperatures. Plasmid preparation and PCR product purification was performed using kits from Macherey Nagel according to protocol. Chemically competent *E. coli* DH5α cells (Thermo Fisher Scientific) were used as a chassis for all cloning approaches and the constructs verified through sequencing (Microsynth).

### Construction of expression plasmids

Plasmids containing LD(4), (3)Mod1(2) and (3)Mod2(2), (3)Mod3(4), DEBS2 and DEBS3 were described previously^27,28,37^.

### Generating KS3 variants

KS3 mutations were introduced by amplifying fragments using the corresponding mutagenic primers (Table 2) with flanking primers S1_fw and S2_rev and pET28b_(3)Mod3(4) as template. Fragments were subsequently cloned into pET28b_(3)Mod3(4) following backbone digest with NcoI and BsiWI. The catalytic knockout of AT3 (Ser651Ala) was constructed similarly, using AT3_KO as flanking primers and restriction enzymes AscI and MauBI for backbone digest. Implementation of those variants into pET28b_DEBS2 was carried out via T4 ligation after insert and backbone digest using restriction enzymes NcoI and BsiWI.

### Protein purification

Expression plasmids containing all DEBS proteins were individually transformed into *E. coli* BAP1^38^ and inoculated in 1 L of TB from a 10 ml LB preculture. After growth of the expression culture to an OD_600_ of approx.0.7, cultures were cooled down to 21°C, induced with a final concentration of 150 µM IPTG and incubated over night at the same temperature. Cells producing either DEBS protein (except LD(4)) were harvested at 5,000 g and resuspended in buffer A, lysed by sonication and centrifuged at 55,000 g at 4°C for 45 min. The supernatant was incubated with preequilibrated Protino Ni-NTA beads (1 ml resin, Macherey-Nagel) at 4°C for 1h. The beads were collected through centrifugation at 500 g at 4°C for 5 min, the supernatant discarded and the beads transferred onto a Protino 14 ml gravity column (Macherey-Nagel). After 20 ml of additional equilibration using buffer A, beads were washed using 15 ml 5 % (v/v) buffer B in buffer A. For elution 8 ml buffer B were used, the eluate filled up to 15 ml with buffer F, loaded onto a 5 ml HiTrap Q anion exchange column (Cytiva) preequilibrated with buffer C and eluted in an 80 ml gradient from buffer C to buffer D. Cells producing LD(4) were processed identically, but only buffer A was used from resuspension to purification over a preequilibrated 1 ml StrepTrap column (Cytiva), supplemented with 2 mM d-desthiobiotin for elution.

Production and purification additional proteins (MatB, RevS, Acx4, CcCcrPAG, Epi)^27,39,40^ were carried out identically, except using *E. coli* BL21(DE3) AI (Thermo Fisher Scientific) and buffer E and F instead of buffer A and B. respectively. Furthermore, no anion exchange was performed, but the proteins desalted on a preequilibrated PD-10 column (Cytiva) using buffer G.

Each protein-containing fractions were pooled and concentrated via an Amicon centrifugal filter using the appropriate molecular weight cutoff and verified via SDS-PAGE. Protein concentration was determined spectroscopically at 280 nm using calculated molar extinction coefficients^41^. The FAD concentration in Acx4 samples was measured at 450 nm and FAD added to reach equimolar protein/FAD concentrations.

### Synthesis and HPLC purification of extender units

Extender unit synthesis was carried out as previously described^27,39^, with minor adoptions. In brief, methylmalonyl-CoA was synthesized using the respective dicarboxylic acid as substrates for MatB. 40 mg CoA (1 eq.), 30.4 mg methylmalonic acid (5 eq.) and 140.8 mg ATP (4 eq.) were dissolved in 8 mL of 200 mM KHCO_3_ containing 15 mM MgCl_2_ and 5 µM MatB. The reaction was incubated at 30°C and 200 rpm, followed until completion using DTNB and quenched with a final concentration of 10 % (v/v) formic acid.

All other extender units (butyl-, 3-methylbutyl-, and hexylmalonyl-CoA) were synthesized in an assay volume of 6 ml, which contained 100 mM HEPES pH = 7.5, 20 mM MgCl2, 100 mM KHCO3, 4.35 mM (20 mg) CoA, 17.38 mM ATP and 20.86 mM of the corresponding carboxylic acid. Upon equilibration of the assay mixture at 30 °C, the enzymes were added to a final concentration of 5 µM ligase (RevS), 3 µM Acx4 and 3 µM CcCcrPAG. The reactions were incubated for 2 h 30°C at 200 rpm, while monitoring CoA consumption using Ellman’s reagent. Upon completion, all reactions were quenched with a final concentration of 10% (v/v) formic acid, centrifuged for 10 min at 5000 g, filtered through a 0.2 µm syringe filter and flash frozen in N_2_ (l) and stored at -80°C if not immediately subjected to HPLC-MS purification.

All malonyl-CoAs were purified via reverse phase LC/MS using a Gemini 10 μm NX-C18 110 Å, 100 × 21.2 mm, AXIA packed column (Phenomenex). Using 50 mM NH4HCO2 pH 4.1 as aqueous phase, the column was equilibrated after injection for 2 min with 5 % MeOH, followed by a gradient from 5 % to 40 % MeOH in 19 min, a 2 min washing step at 95 % MeOH and a re-equilibration step of 3 min at 5 % MeOH. The flow rate was kept constant at 25 ml min^-1^. Fractions containing the product were pooled, flash frozen in liquid N2, lyophilized and stored at -20°C until use. The concentration was determined via UV/Vis at 260 nm using an exctinction coefficient ε_CoA 260 nm_ = 16.4 mM^-1^ cm^-1^.

### Extender unit incorporation assays and assay optimization

The final in vitro assay setup contained 200 mM sodium phosphate buffer (pH 7.2), 2 mM NADPH, 4 µM methylmalonyl-CoA epimerase, 0.8 mM propionyl-CoA, 2 mM methylmalonyl-CoA and 1 mM of either non-natural alkylmalonyl-CoA in a final volume of 50 µM. All DEBS proteins were added in equimolar concentrations (2 µM) with the exception of (3)Mod2(2), which was added in twofold excess (4 µM).

Samples were taken after 5 hours and 17 hours, quenched with a final concentration of 10 % (v/v) formic acid, centrifuged at 17,000 g for 10 min and either stored at -80°C or directly subjected to HPLC-TOF measurement.

### HPLC measurements of polyketides

LC-high resolution MS analysis was carried out using an Agilent 6550 iFunnel Q-TOF LC-MS system equipped with an electrospray ionization source set to positive ionization mode.

Polyketides were separated on a Zorbax SB-C18 column (50 mm x 2.1 mm, particle size 1.8 µm, Agilent) using water (A) and acetonitrile (B) enriched with formic acid to a final concetration of 0.1%. The gradient condition is as follows: 0 min 5% B; 1 min 5% B; 6 min 95% B, 6.5 min 95% B, 7 min 5% B at a flow rate of 250 µl/min. Capillary voltage was set at 3.5 kV and nitrogen gas was used as nebulizing (20 psig), drying (13 l/min, 225 °C) and sheath gas (12 l/min, 40°C). MS data were acquired with a scan range of 100-1000 m/z.

Data was analyzed using MassHunter Qualitative Analysis software and MassHunter TOF Quantitative Analysis (Agilent) using the m/z ratios given in Table S1.

### Structural elucidation of products via LC MS-MS

LC-high resolution MS-MS analysis was carried out using Thermo Scientific Vanquish HPLC System coupled to a Thermo Scientific Orbitrap ID-X with electrospray ionization source set to positive ionization mode. The tetraketides isomers were separated on a Zorbax SB-C18 column (50 mm x 2.1 mm, particle size 1.8 µm, Agilent) using water (A) and acetonitrile (B) enriched with formic acid to a final concentration of 0.1%. The gradient condition is as follows: 0 min 5% B; 3 min 5% B; 18 min 95% B, 21 min 95% B, 21.1 min 5% B, 25 min 5% B with a flow rate of 250 µl/min.

A targeted MS2 method was used for structural elucidation. The full scan measurements were conducted applying an Orbitrap mass resolution of 60 000 without quadrupole isolation in a mass range of 150-400 in profile mode. The targeted masses of the tetraketides were isolated using a quadrupole isolation window of

0.4 *m/z*. Collision induced dissociation was performed in the ion routing multipole with a relative collision energy of 30 %. Fragments were detected using the Orbitrap at a predefined mass resolution of 30 000 in the range between 50 and 300. The three targeted masses were 271.1904 (**2b²** and **2b³**), 285.206 (**2c²** and **2c³**), and 299.2217 (**2d²** and **2d³**).

Compound Discoverer 3.2 (CD 3.2, Thermo Fisher Scientific) was used for the spectra analysis and acquiring the possible fragment structure from each tetraketide. The pre-defined workflow “E and L Expected w FISh Scoring” was used for the analysis after removing the nodes “Create Analog Trace” and “Mark Background Compounds” (Figure S9). The structures of **2b²-d²** and **2b³-d³** were included in the “Expected Compounds” Library and in the node “Generate Expected Compounds”. The “FISh Scoring” node contains the *in silico* fragmentation algorithm which will match the theoretical fragments (Figure S9) to the MS/MS spectra.

Differences in the MS/MS spectra of all isomer groups (**2b² & 2b³, 2c² & 2c³** and **2d² & 2d³**) could be attributed to the changes of fragment intensity between the loss of one water molecule (base peak of the MS/MS spectrum) or two water molecules. The loss of the different number of water molecules were the main fragments of the MS/MS spectra and it was not possible to define which pattern belonged to which isomer. In the MS/MS spectra of the peak corresponding to the isomer that was eluting first (Figure S10A, S11A, S12A), the 2xH_2_O water loss fragment was higher than 30 % corresponding to the base peak for the different masses of each isomer pair, while it was below 25% from all the MS/MS spectra of the isomer eluting second (Figure S10B, S11B, S12B). The fragment 127.1116 (C8H15O) was the only fragment which was higher than the 10% from the base peak and present in consistence in one of the two isomer for all of the groups. Possible mechanisms to create the detected fragment (Figure S10C, S11C, S12C) show a much higher probability of compounds **2b³, 2c³** and **2d³** to create this, whereas the fragmentation mechanism for their structural isomers would be significantly more complicated. Thus, we concluded that the first elution peak would correspond to **2b³, 2c³** and **2d³**, while the second peak corresponds to **2b², 2c²** and **2d²**.

## Notes

### Competing Interest Statement

The authors have declared no competing interest.

