## Supplementary Information for "Ketosynthase engineering enhances activity and shifts specificity towards non-native extender units in type I linear polyketide synthase"

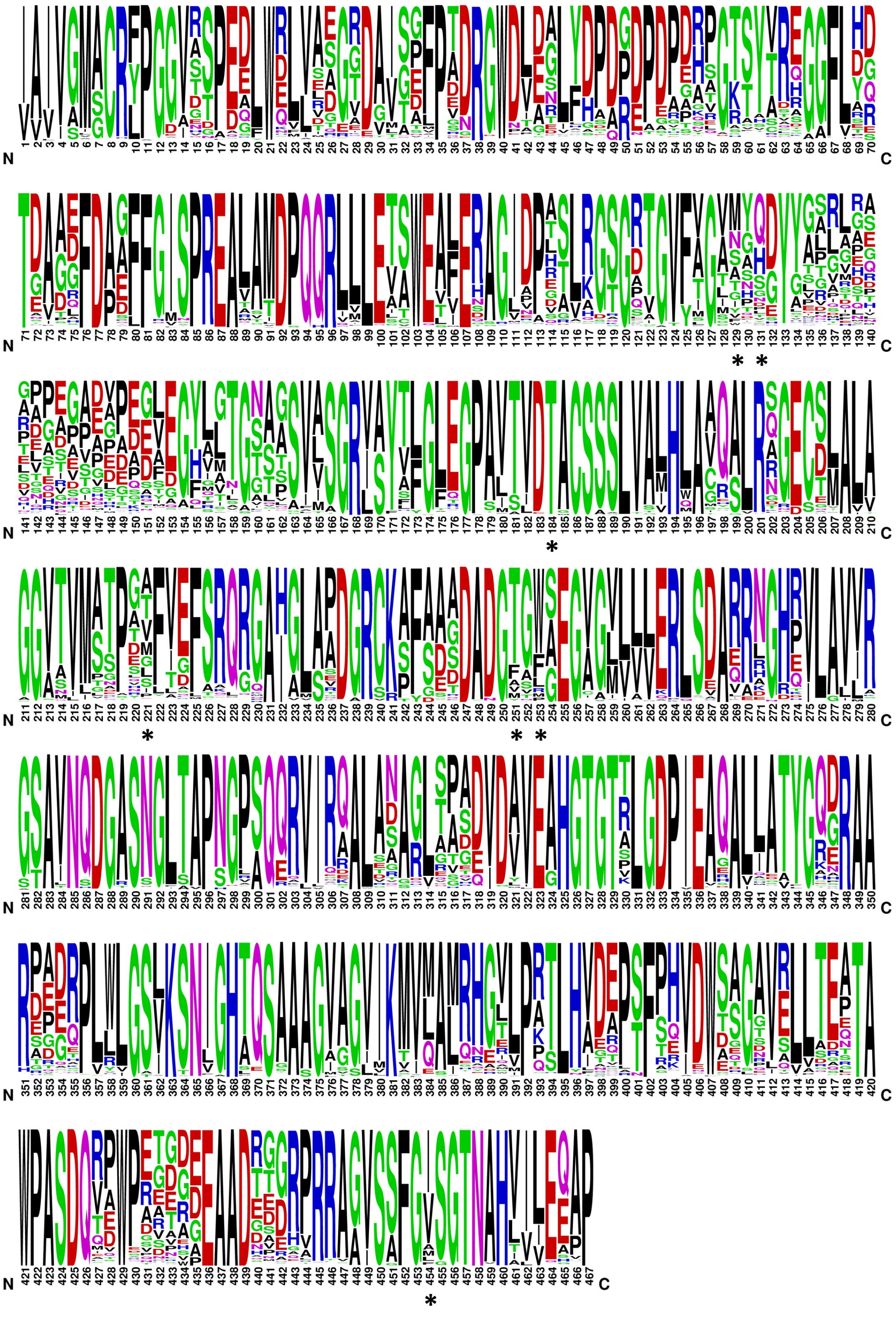

**Figure S1. Sequence frequency of all verified type I KS domains**. Sequence data was retrieved from ClusterCAD[1] and the sequence logo created using the WebLogo online tool[2,3]. Note that the numbering diverges from that used throughout this work. Residues correspond to the following numbers: A154 = 129, F156 = 131, T200 = 184, V237 = 221, F263 = 251, F265 = 253, I444 = 454 and are indicated by asterisks.

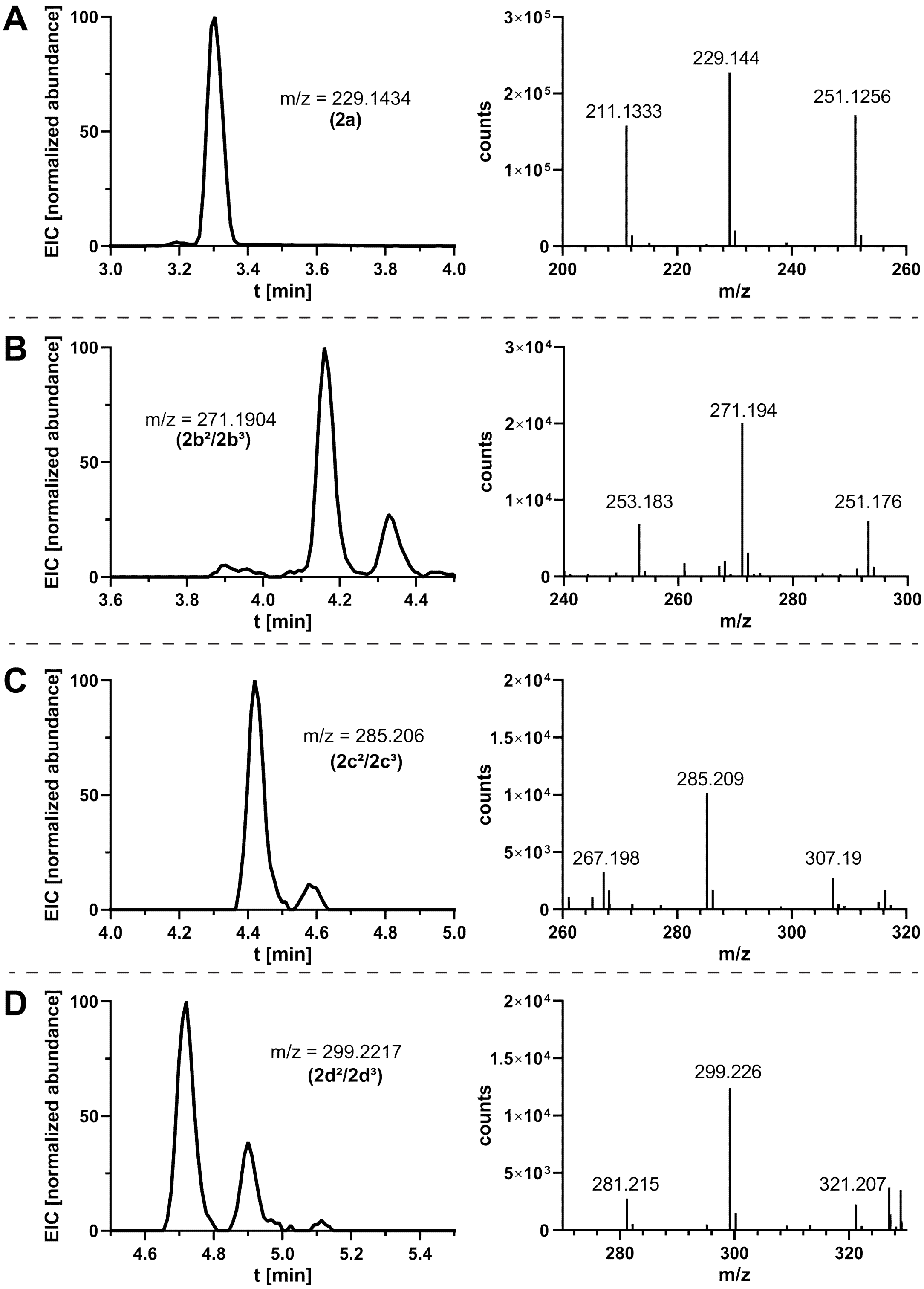


**Figure S2. Extracted ion chromatograms, expected m/z and measured MS spectra for compounds 2a, $2b^2$-$d^2$ and $2b^3$-$d^3$.** Measurements for all chromatograms stem from representative assay samples. Shown are compounds 2a (methyl-tetraketide, **A**), $2b^2$/$2b^3$ (butyl-tetraketide, **B**), $2c^2$/$2c^3$ (3-methylbutyl-tetraketide, **C**) and $2d^2$/$2d^3$ (hexyl-tetraketide, **D**). m/z is given as [M+H]+. Determination of each structural isomer is shown in Figure S10-S12.

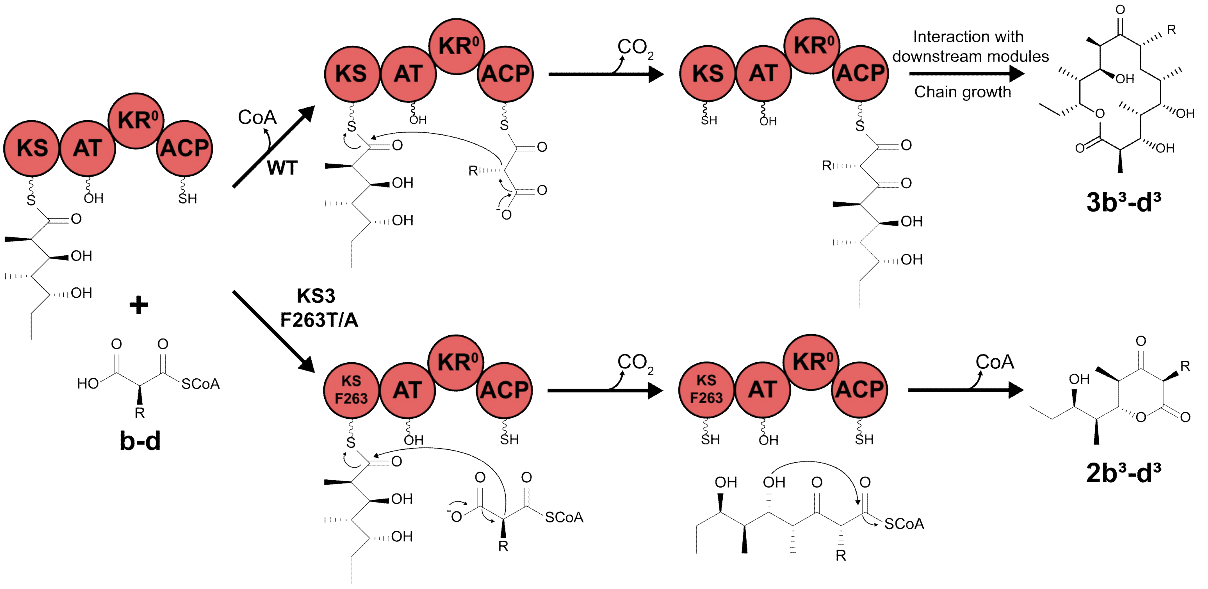


**Figure S3. Canonical mechanism of chain elongation and proposed shunting mechanism.** In the wildtype system (upper), the AT domain selects for an extender unit and transfers it onto the ACP. The malonyl-ACP enters the KS active site and forms an enolate through decarboxylation, which enables chain elongation via condensation. The elongated polyketide chain remains tethered to the ACP until transacylation to the downstream KS and further processing. In the F263 shunting mechanism (lower), the opened KS active site allows for the malonyl-CoA extender unit to act as the nucleophile. While following the identical mechanism, the intermediate remains CoA-tethered and will eventually undergo intramolecular cyclization, resulting in tetraketide shunt product **2b³-d³**.

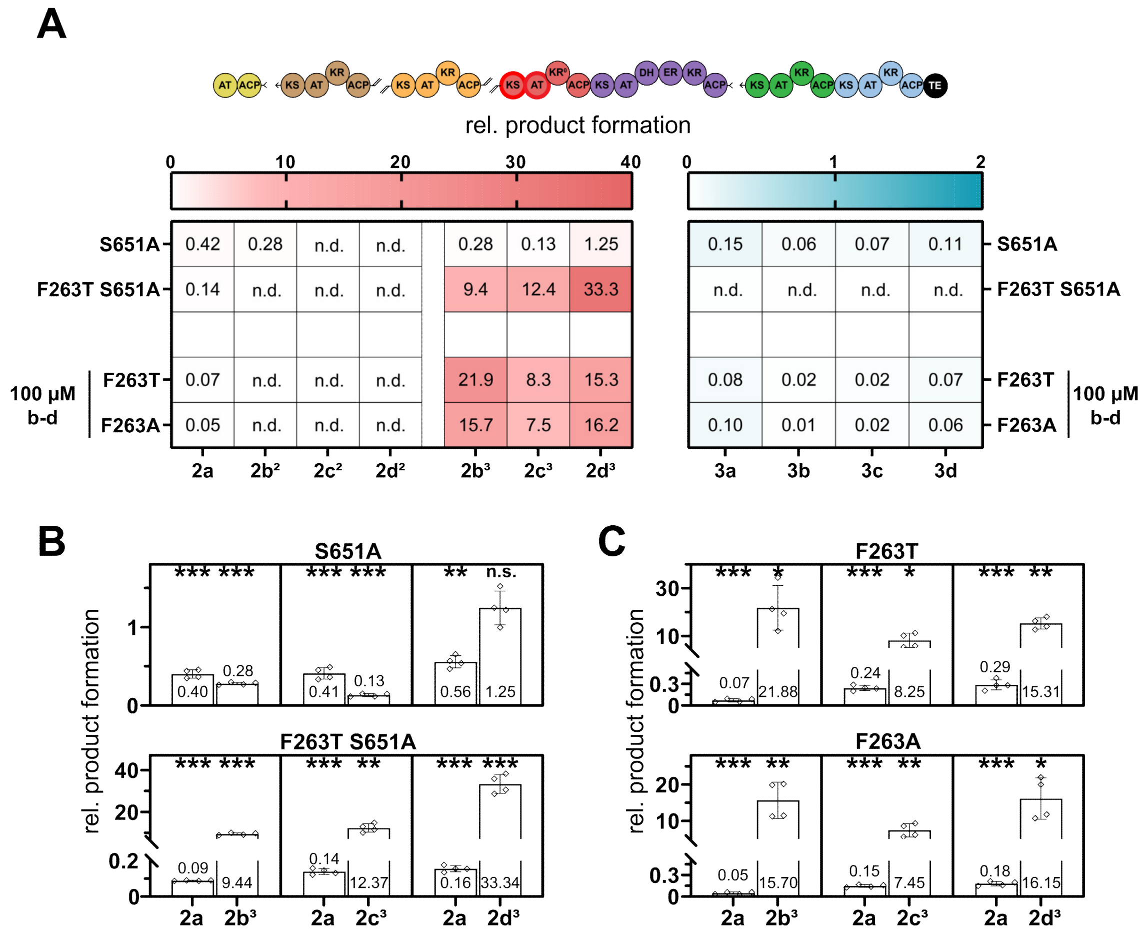


**Figure S4. Relative product formation trends and product distribution of the catalytic AT3 knockout tested in the full assembly line and F263 variants using 100 µM non-native extender unit.** (**A**) Product formation of all tetraketides, as well as 6-dEB and analogues was severely impeded when mutating the catalytic S651 to Ala, but residual activity remained. When combined with the module 3 F263T variant, product formation greatly resembled the proposed tetraketide shunt observed for the F263 variants alone, hinting towards an AT3-independent shunting mechanism. Full assembly line activity is not restored when using a lower concentration of **b-d**. Product distribution is overall decreased by the AT3 knockout, but F263T restored the specificity shift towards the non-native extender unit (**B**). An identical trend can be observed when adding lower (i.e. non-competitive) amounts of **b-d** to either F263 variant (**C**), further highlighting the probability of an AT-ACP bypassing mechanism. Values are derived from the detected EIC of each compound compared to that of the wild type after 17 h and display the mean of n = 2 technical replicates. Significance ($H_0$: relative product formation = 1) was determined using a two-tailed one-sample t-test. *$p \leq 0.05$, **$p \leq 0.005$, ***$p \leq 0.0005$. n.d. – not detected. n.s. – not significant.

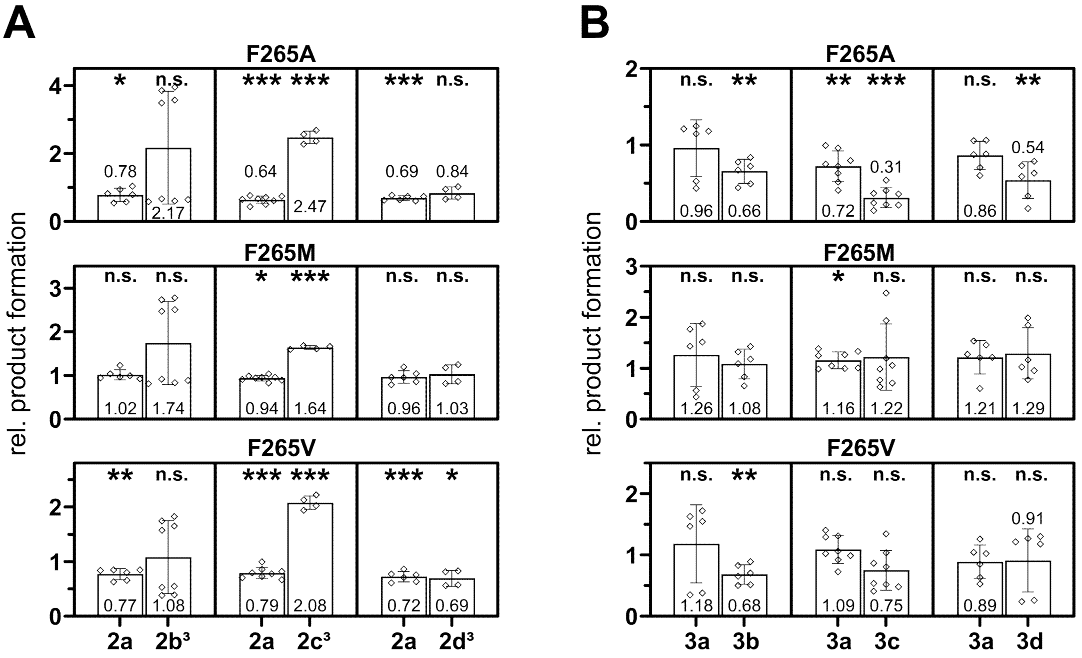


**Figure S5. Product distribution of F265 variants in the full assembly line (related to Figure. 3).** Displayed is the relative product formation by all F265 variants for 2a and either non-natural tetraketide (**A**) and 6-dEB and analogues (**B**). Values are derived from the detected EIC of each compound compared to that of the wild type after 17 h and display the mean of n = 2 biological replicates (n = 2 technical replicates each). Significance ($H_0$: relative product formation = 1) was determined using a two-tailed one-sample t-test. *$p \leq 0.05$, **$p \leq 0.005$, ***$p \leq 0.0005$. n.s. – not significant.

**Combining AT2 and KS3 variants**

Given the unexpected, yet significant impact of some KS3 variants on product formation, we hypothesized them to "de-bottleneck" shunt product accumulations we had observed from testing previously described AT2 variants[4-6]. Those had displayed a significantly enhanced promiscuity towards **b-d** with up to 10-fold increases of product formation in a truncated triketide system (terminated after module 2, Figure 6A). Transfer of those variants into the full DEBS assembly line partially resulted in a similar accumulation of hop-off shunt product **1b-d**, whereas formation of 6dEB and analogues **3a-d** was severely reduced (Figure 6B).

To test this possibility, we combined the KS3 variants with the AT2 variants in both the system terminated after module 3 as well as in the full assembly line. Should the downstream limitation arise from the ketosynthase any variant display a higher promiscuity for derivatized intermediates, a combination should increase the product formation of any given product downstream of module 2. The M3 F263 variants were omitted from this approach, since their impact overlaid all effects of other substitutions.

Here, the KS3 variants showed only a minor effect of increased tolerance towards a module 2 derived non-native intermediate (Figure S7), with mostly insignificant or < 3-fold increases of **2b$^2$-d$^2$** tetraketide. Furthermore, many AT2-KS3 variant combinations led to an increased accumulation of triketide compared to the AT2 variants, indicating a decrease in promiscuity towards an intermediate carrying a C2-substitution. The only increase observed was from module 3 derived tetraketide **2b$^3$-d$^3$** in an identical fashion to the KS3 variants alone (Figure 2), with F265 variants displaying the most prominent increase of up to 20-fold.

We nevertheless also tested any variant combinations, which displayed a ≥ 2-fold increase for **2b$^2$-d$^2$** in the full assembly line (Figure S8). Here the only significant positive increase was from **1b-d** accumulation using the Y754V variant, and any combination containing a KS3 F265 variant, in which accumulation of **2d$^3$** was increased. No significant increases (or specificity changes, data not shown) regarding the final polyketide **3a** or any of its analogues **3b-d** could be observed. It must be noted, however, that the effects of the AT2 variants themselves had been far less prominent than those of the KS3 variants had, and as much might have not sufficed to produce detectable changes.

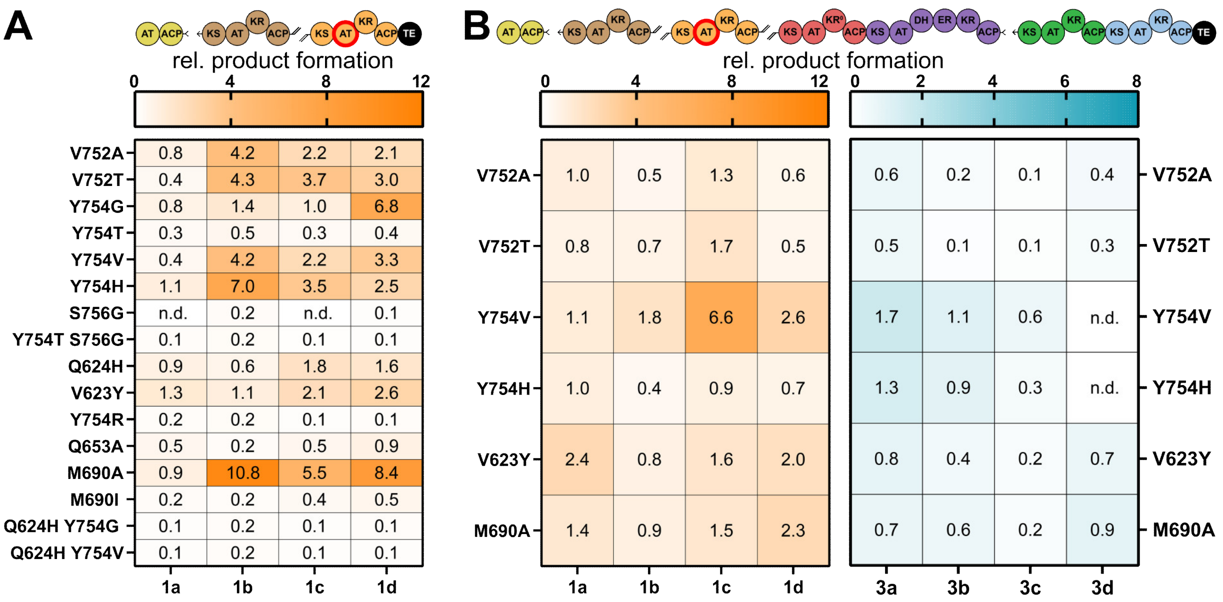


**Figure S6. Relative product formation trends of AT2 variants.** AT2 variants were initially tested in the triketide system (**A**), which terminates after module 2 through C-terminal fusion of the DEBS TE domain[7]. Transfer of the same mutations into module 2 resulted in largely insoluble enzyme variants. Those which could be purified were tested in the full DEBS assembly line in identical conditions (**B**). While 6-dEB formation decreased or remained stagnant, the significant accumulation of module 2 derived triketide **2b-d** prevailed for some variants, albeit at lower yields. Color usage according to Figure 1. Values are derived from the detected EIC of each compound compared to that of the wild type after 17 h and display the mean of n = 2 technical replicates. n.d. not detected.

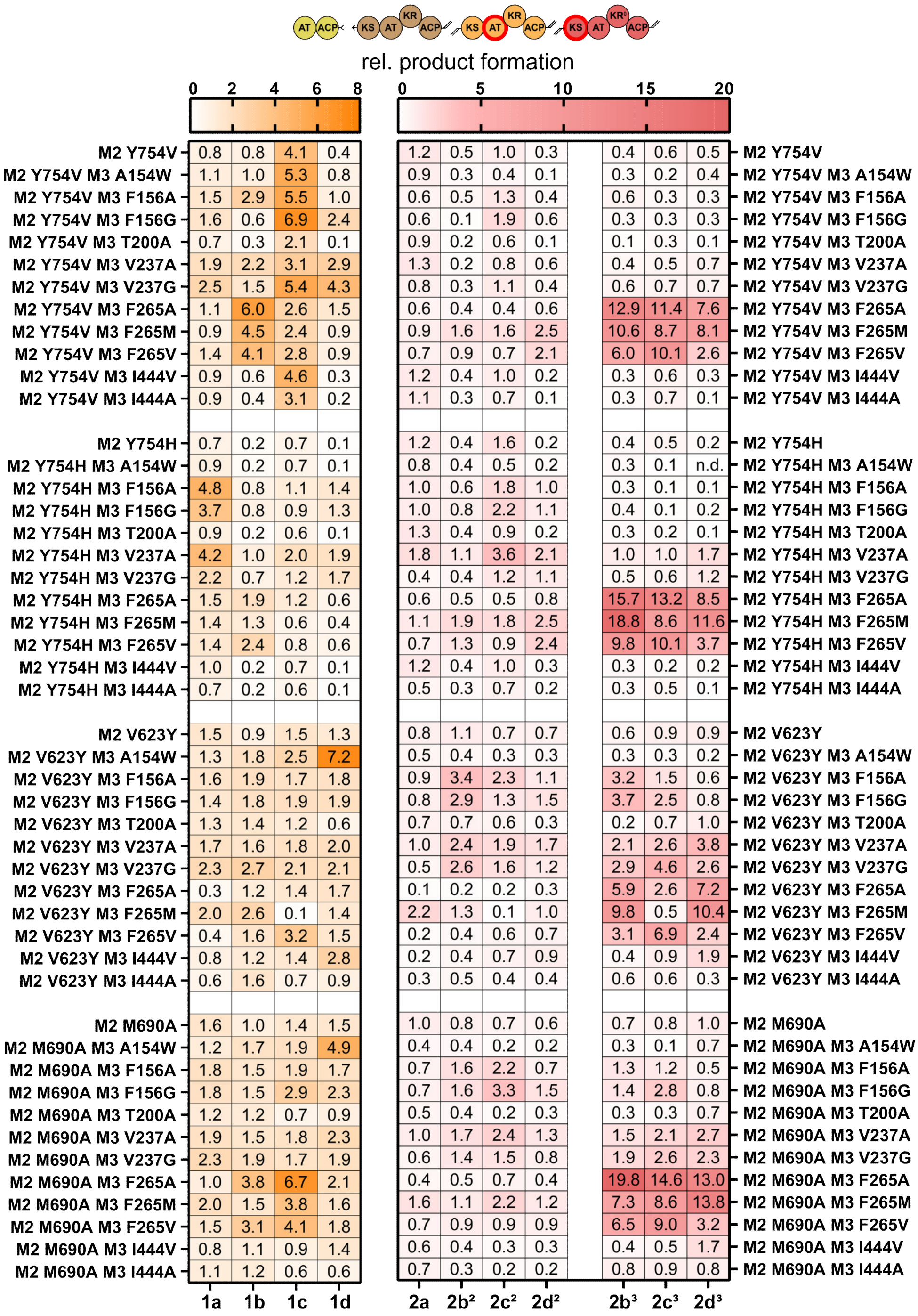


**Figure S7. Relative product formation trends of AT2-KS3 variant combinations in the tetraketide system.** The AT2 variants are shown on top of each set followed AT2-KS3 combinations. Displayed are the relative formation of triketide, tetraketide derived from incorporation at module 2, and tetraketide derived from incorporation at module 3 (**2b**$^3$-**d**$^3$). Those variant combinations, which displayed at least a twofold increase of either **2b-d** were further tested in the full assembly line (Figure S8). Color usage according to Figure 1. Values are derived from the detected EIC of each compound compared to that of the wild type after 17 h and display the mean of n = 2 technical replicates. n.d. – not detected.

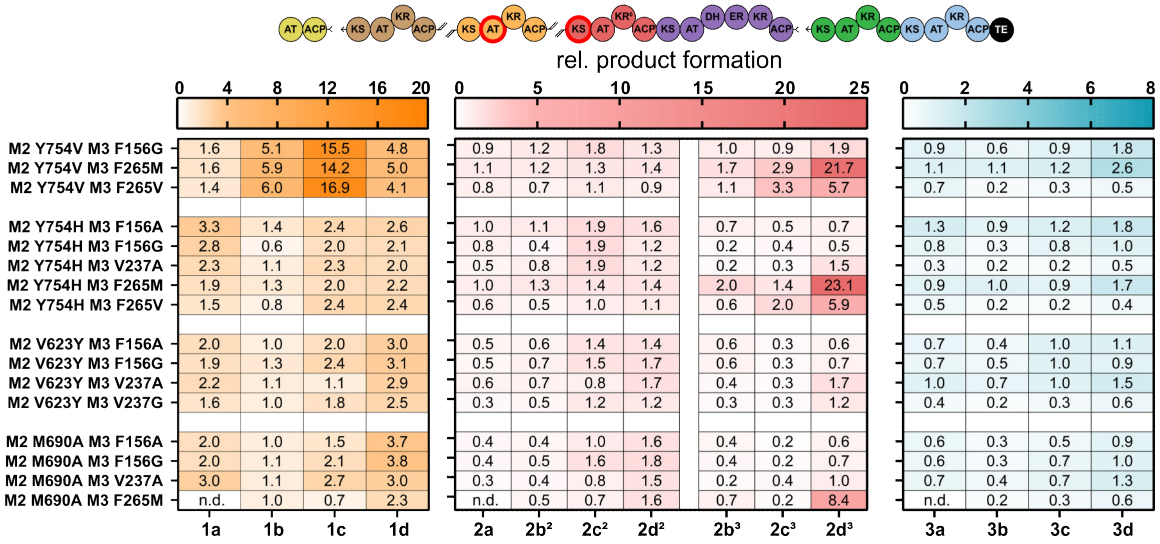


**Figure S8. Relative product formation trends of selected AT2-KS3 variant combinations in full assembly line.** No significant increases of any 6-dEB analogue could be detected. While the F265M variant had no clear effect in the full assembly line on its own (Figure 3), combinations with AT2 variants led to an increase of d derived tetraketide shunt product **2d³**. Color usage according to Figure 1. Values are derived from the detected EIC of each compound compared to that of the wild type after 17 h and display the mean of n = 2 technical replicates. n.d. not detected.

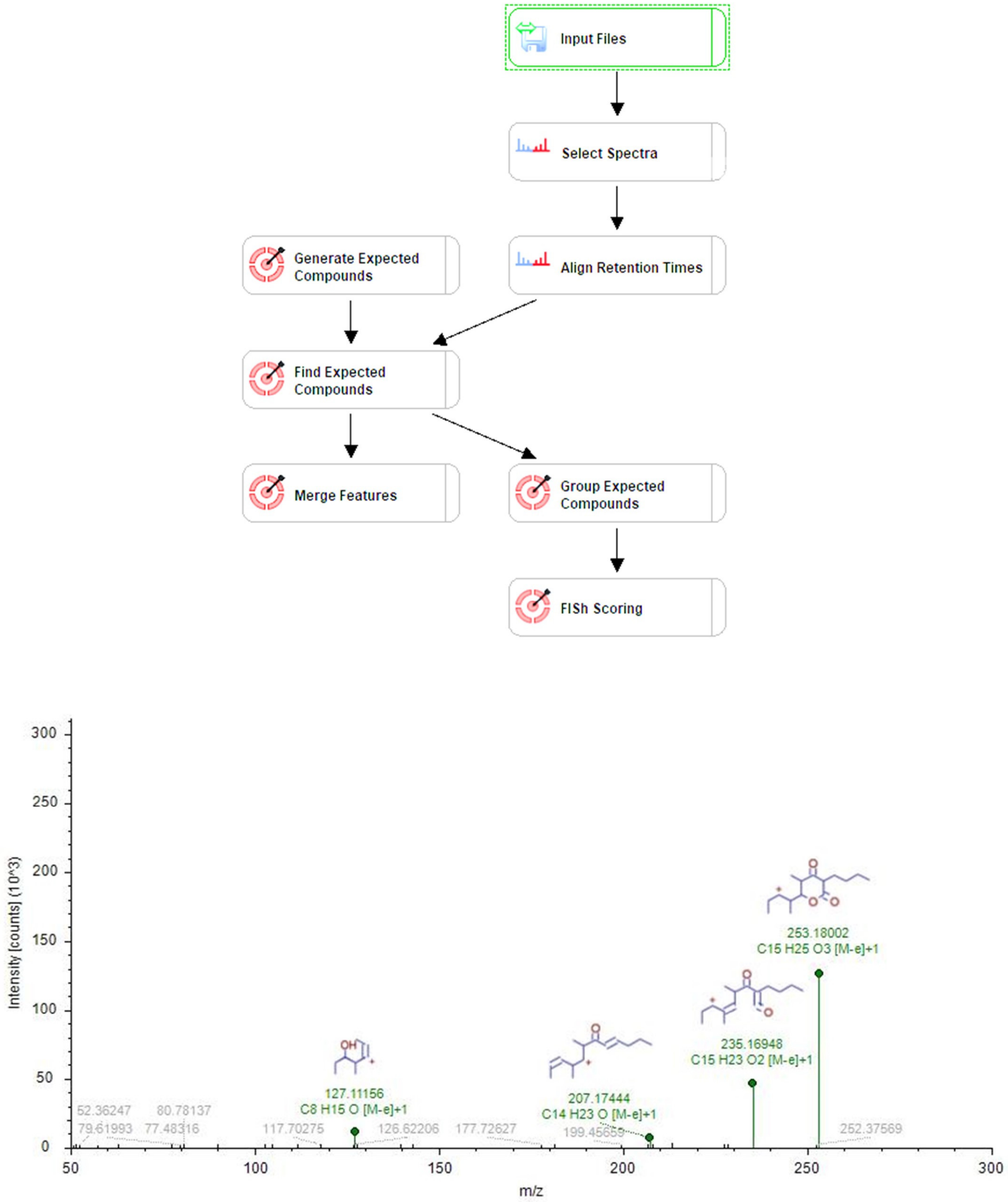


**Figure S9. Compound Discoverer 3.2 workflow.** Proposed structures by CD 3.2 MS/MS for compound 2b* of all theoretical fragments found in the spectrum.

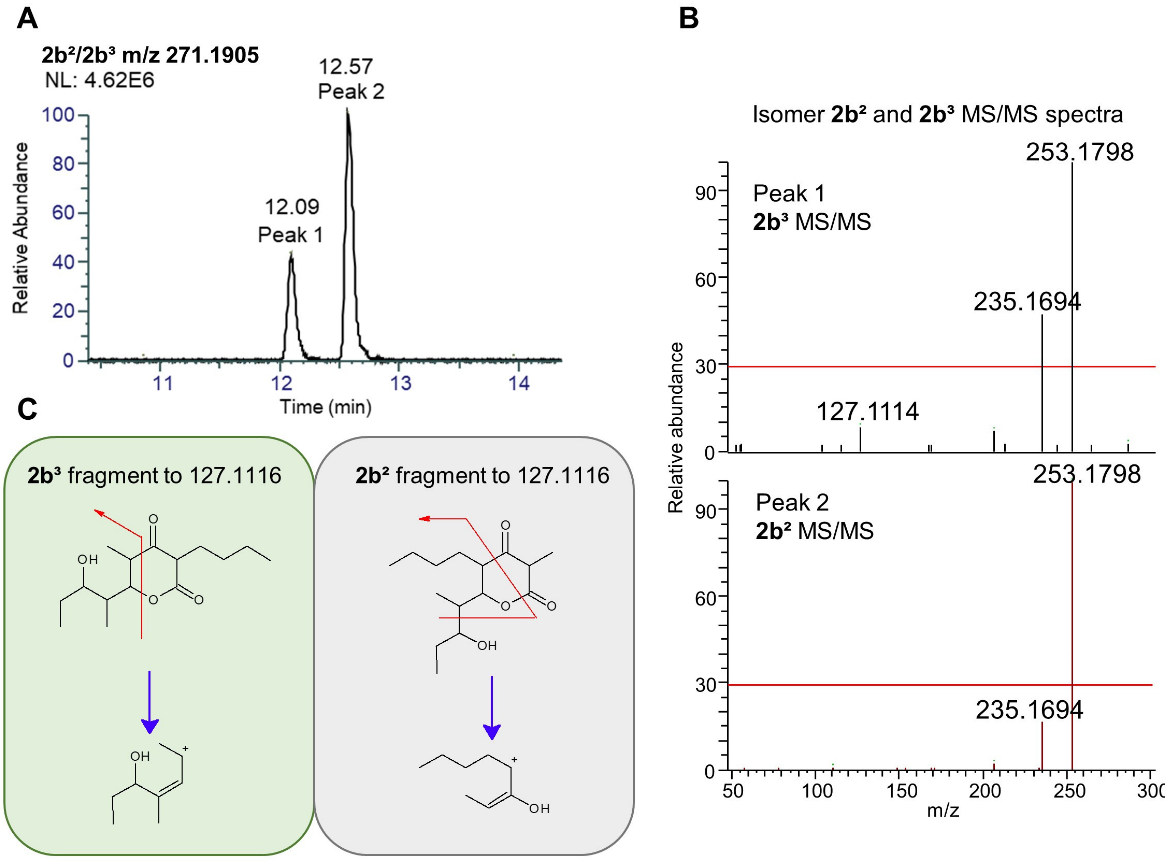


**Figure S10. Structural elucidation of compounds 2c and 2c*.** The extracted ion chromatogram of the two compounds (**A**) and the MS/MS spectra from the two different peaks and their differences (**B**), according to which the first peak corresponds to compound $\mathbf{2b^3}$ and the second peak to compound $\mathbf{2b^2}$. (**C**) the proposed fragmentation to 127.1116 ($C_8H_{15}O$) according to CD 3.2.

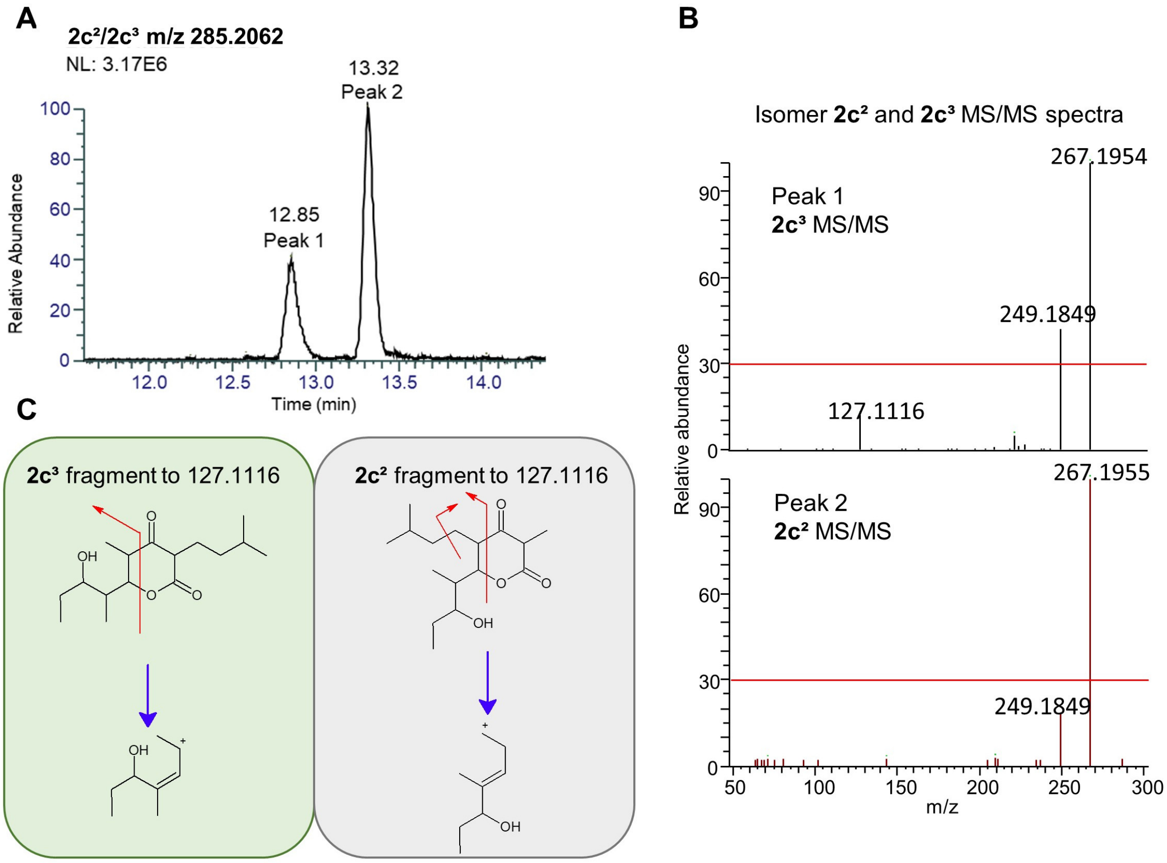


**Figure S11. Structural elucidation of compounds 2c and 2c*.** The extracted ion chromatogram of the two compounds (**A**) and the MS/MS spectra from the two different peaks and their differences (**B**), according to which the first peak corresponds to compound **2c$^3$** and the second peak to compound **2c$^2$**. (**C**) the proposed fragmentation to 127.1116 ($C_8H_{15}O$) according to CD 3.2.

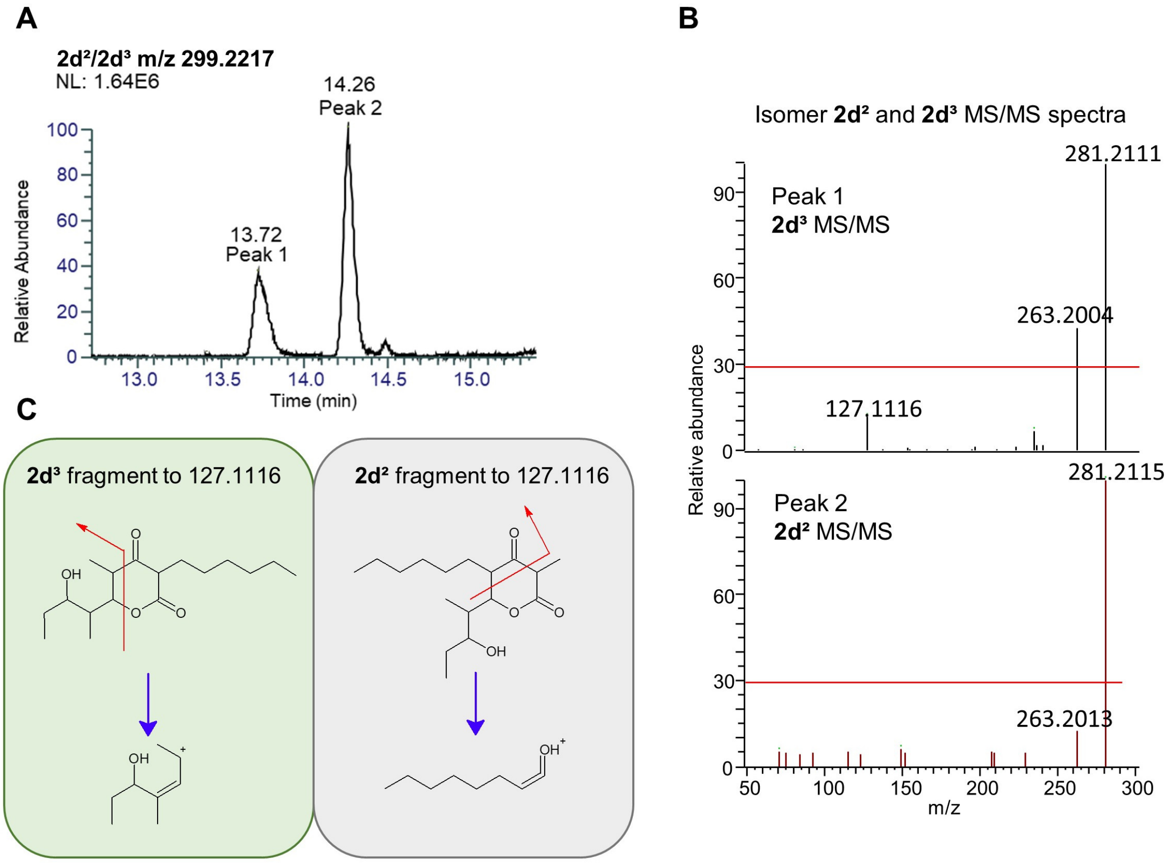


**Figure S12. Structural elucidation of compounds 2d$^2$ and 2d$^3$.** The extracted ion chromatogram of the two compounds (**A**) and the MS/MS spectra from the two different peaks and their differences (**B**), according to which the first peak corresponds to compound **2d$^3$** and the second peak to compound **2d$^2$**. (**C**) the proposed fragmentation to 127.1116 ($C_8H_{15}O$) according to CD 3.2.

**Table S1. Overview of mass spectrometry adducts detected for product analysis**.

| Compound | | **1a** | **1b** | **1c** | **1d** |
|---|---|---|---|---|---|
| MS adduct | [M+H]+ | 171.1016 | 213.1485 | 227.1642 | 241.1798 |
| | [M-H2O+H]+ | 153.090457 | 195.137407 | 209.153057 | 223.168707 |
| | [M+Na]+ | n.d. | n.d. | n.d. | n.d. |
| Compound | | **2a** | **$2b^2/b^3$** | **$2c^2$/c 3** | **$2d^2/d^3$** |
| MS adduct | [M+H]+ | 229.143436 | 271.190386 | 285.206036 | 299.221686 |
| | [M-H2O+H]+ | 211.132322 | 253.179272 | 267.194922 | 281.210572 |
| | [M+Na]+ | 251.124832 | 293.171782 | 307.187432 | 321.203082 |
| Compound | | **3a** | **3b** | **3c** | **3d** |
| MS adduct | [M+H]+ | 387.2741 | 429.3211 | 443.3367 | 457.3524 |
| | [M-H2O+H]+ | 369.263001 | 411.309952 | 425.325602 | 439.341252 |
| | [M+Na]+ | 409.2566 | 451.3056 | 465.3192 | 479.3349 |
